# Distinct Representations of Body and Head motion are Dynamically Encoded by Purkinje cell Populations in the Macaque Cerebellum

**DOI:** 10.1101/2021.10.25.465748

**Authors:** Omid A. Zobeiri, Kathleen E. Cullen

## Abstract

The ability to accurately control our posture and perceive spatial orientation during self-motion requires knowledge of the motion of both the head and body. However, whereas the vestibular sensors and nuclei directly encode head motion, no sensors directly encode body motion. Instead, the integration of vestibular and neck proprioceptive inputs is necessary to transform vestibular information into the body-centric reference frame required for postural control. The anterior vermis of the cerebellum is thought to play a key role in this transformation, yet how its Purkinje cells integrate these inputs or what information they dynamically encode during self-motion remains unknown. Here we recorded the activity of individual anterior vermis Purkinje cells in alert monkeys during passively applied whole-body, body-under-head, and head-on-body rotations. Most neurons dynamically encoded an intermediate representation of self-motion between head and body motion. Notably, these neurons responded to both vestibular and neck proprioceptive stimulation and showed considerable heterogeneity in their response dynamics. Furthermore, their vestibular responses demonstrated tuning in response to changes in head-on-body position. In contrast, a small remaining percentage of neurons sensitive only to vestibular stimulation unambiguously encoded head-in-space motion across conditions. Using a simple population model, we establish that combining responses from 40 Purkinje cells can explain the responses of their target neurons in deep cerebellar nuclei across all self-motion conditions. We propose that the observed heterogeneity in Purkinje cells underlies the cerebellum’s capacity to compute the dynamic representation of body motion required to ensure accurate postural control and perceptual stability in our daily lives.

## Introduction

The cerebellum guides motor performance by computing differences between the expected versus actual consequences of movements and then adjusting the commands sent to the motor system (reviewed in: Wolpert et al., 1998, Raymond and Medina 2018). Patients with damage to the anterior vermis of the cerebellum show impaired posture and balance, as well as deficits in motor coordination (Dichgans and Diener, 1984; Bastian et al., 1998; Giese et al., 2008; Sullivan et al., 2005; Mitoma et al., 2021). In this context, the anterior vermis has a vital role in the vestibulospinal pathways that generate the postural adjustments required to ensure the maintenance of balance during our everyday activities. Additionally, there is an emerging consensus that the cerebellum contributes to our self-motion perception. Indeed, patients with degeneration of the cerebellar vermis demonstrate reduced perceptual time constants and detection thresholds to externally applied rotations (Bronstein et al. 2008; Dahlem et al., 2016).

Inherent to the generation of vestibulospinal reflexes and stable perception of our motion is the requirement that central vestibular pathways explicitly transform vestibular information from a head-centered to a body-centered reference frame. This is because the vestibular sensory organs are located within the head, making the vestibular system’s native reference frame head-centered (reviewed in Cullen 2019). In turn, vestibular nerve afferents and their targets in the vestibular nuclei also encode information in a head-centered reference frame (Roy and Cullen 1998, 2001, 2004; Carriot et al. 2013; Brooks and Cullen 2014; Sadeghi et al., 2007; Jamali et al., 2009; Cullen and Minor 2002). However, the brain must account for the position of the head relative to the body for vestibulospinal reflexes to accurately control the musculature required to maintain upright posture and balance (Tokita et al., 1989, 1991; Kennedy and Inglis 2002). Neck proprioceptors provide this essential head position information (reviewed in Cullen and Zobeiri 2021). Thus, the integration of neck proprioceptive and vestibular signals is thought to underlie the transformation from a head-centered to a body-centered reference frame required in vestibulospinal reflexes pathways as well as our ability to perceive body motion independently of head motion (Mergener et al., 1997; Peterka 2002).

There are many reasons to believe that the anterior region of the cerebellar vermis is vital in the transformation of vestibular information from a head-centered to a body-centered reference frame. First, Purkinje cells in this region project to the rostral portion of the fastigial nucleus, the most medial of deep cerebellar nuclei (Batton et al., 1977; Yamada and Noda 1987), which lesion studies have shown serves an important role in the control of posture and balance (Thach et al., 1992; Kurzan et al., 1993; Pelisson et al., 1998). Second, inhibition of the cerebellar vermis via continuous theta-burst stimulation impairs the modulation of vestibulospinal pathways that normally accounts for changes in the position of the head relative to the body (Lam et al., 2016). Third, neuronal recordings from anterior vermis Purkinje cells in decerebrate cats have demonstrated that individual neurons can encode both vestibular and neck proprioceptive-related information (Manzoni et al. 1998, 1999, 2004), thereby providing a neural substrate for the coordinate transformation using neck proprioceptive signals to convert head-centered vestibular-signals to a body-centered reference frame. However, these studies stopped short of establishing whether Purkinje cells integrate vestibular and neck proprioceptive signals to dynamically encode head or body movement.

Thus, a key question yet to be answered is: Does the cerebellum integrate vestibular and neck proprioceptive signals to provide a dynamic representation of body motion relative to space? Here we recorded the activity of single Purkinje cells in the anterior vermis during head motion, body motion, and combined head and body motion. We found considerable heterogeneity across individual Purkinje cells in their encoding of head versus body motion, with most (~75%) neurons dynamically encoding an intermediate representation of self-motion between head and body motion. These neurons, termed bimodal neurons, responded to both vestibular and neck proprioceptive stimulation and displayed head-position-dependent tuning in their sensitivity to vestibular stimulation. In contrast, a minority of cells, termed unimodal neurons, only responded to vestibular stimulation and unambiguously encoded the motion of the head in space. Across all cells, the linear combination of a given neuron’s response sensitivity to dynamic neck and vestibular stimulation alone well estimated its response during combined stimulation. Finally, we found that a simple linear combination model combining the responses of ~40 Purkinje cells could account for the more homogeneous responses of target neurons in the deep cerebellar nuclei (i.e., the rostral fastigial nucleus) during applied self-motion. Our results provide the first evidence, at the level of single Purkinje cells, that a sequential transformation from a head-centered to body-centered reference frame occurs between the cerebellum and deep cerebellar nucleus to ensure postural and perceptual stability in everyday life.

## Results

### The most vestibular sensitive Purkinje cells in the anterior vermis are also sensitive to stimulation of neck proprioceptors

Each Purkinje cell in our population (n = 73) was responsive to vestibular stimulation generated by whole-body rotation (see Methods) and was insensitive to eye movements. As is illustrated in **Figure 1A**, we found considerable heterogeneity in vestibular sensitivities to ipsilaterally versus contralaterally directed head rotations. Some neurons generated excitatory versus inhibitory responses for oppositely directed head movements (**Fig. 1A**; left, linear). Alternately, some neurons generated bidirectional excitatory responses (center, v-shaped), while others largely only generated excitatory responses for one movement direction (**Fig. 1A**; right, rectifying). This contrasts with the vestibular responses recorded in areas targeted by Purkinje cells in the anterior vermis. Notably, vestibular-only neurons in the rFN and vestibular nucleus consistently show excitatory versus inhibitory responses for oppositely directed head movements (rFN: Gardner and Fuchs, 1975, Shaikh et al., 2005; vestibular nucleus: Scudder and Fuchs, 1992, Cullen and McCrea, 1993, McCrea et al., 1999, Roy and Cullen, 2004).

**Figure 1.**
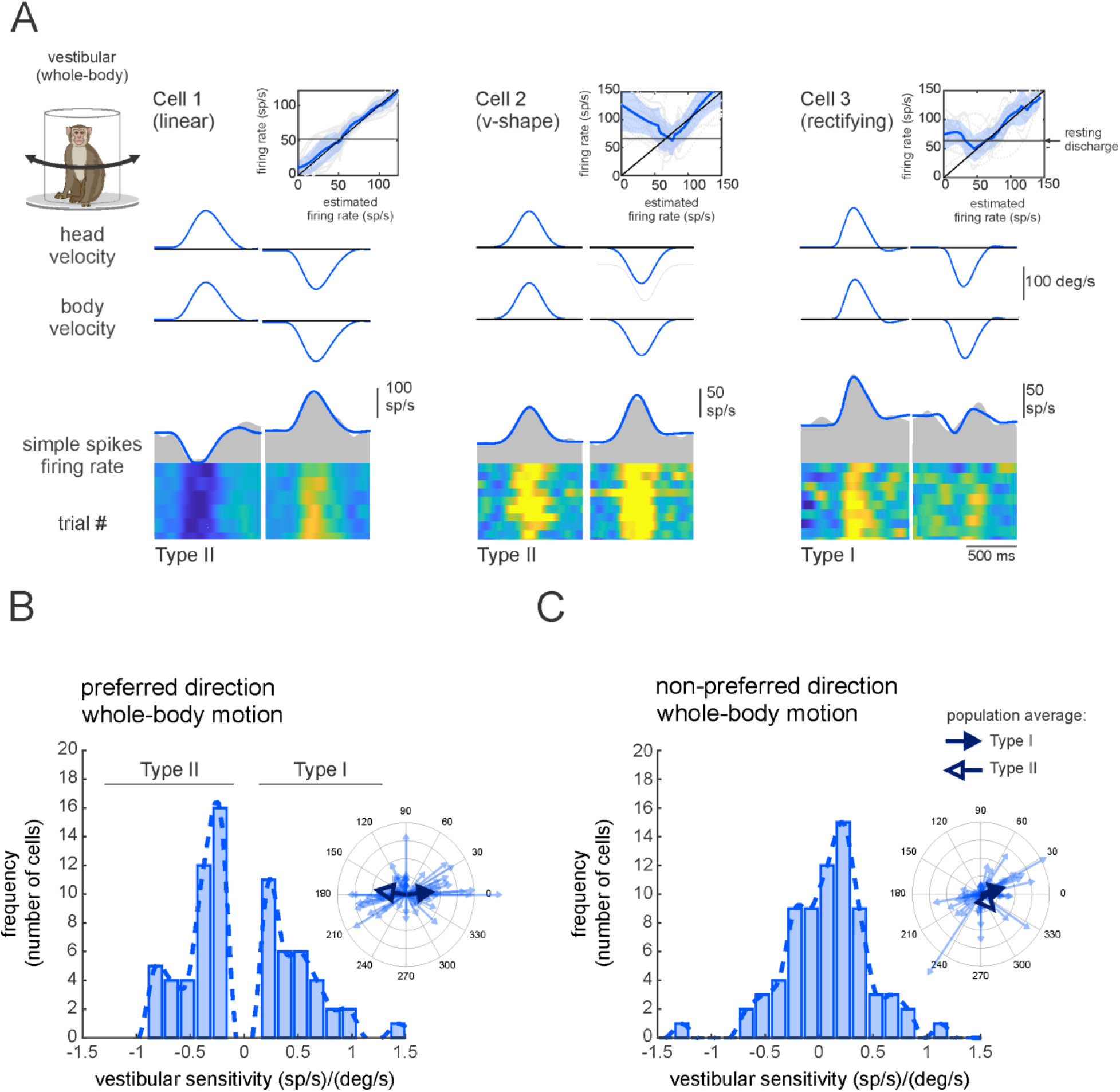
Purkinje cell simple spike responses to vestibular stimulation. **(A)** Vestibular stimulation was generated by applying passive whole-body rotations about the vertical axis (i.e., vestibular stimulation condition). The resulting neural responses are shown for three example Purkinje cells. The top two rows illustrate rotational head versus body velocities. The bottom row shows the simple spike firing rate (gray shaded regions) with the linear estimation of the firing rate based on head motion superimposed (blue traces). *Insets*: the relationship between simple spike firing rate (phase-corrected) and angular head-in-space velocity. **(B,C)** Distribution of vestibular sensitivities for motion in the preferred **(B)** direction (i.e., the direction resulting in the larger increase in simple spike firing rate) and non-preferred **(C)** direction. Note, by convention positive and negative values in **(B)** represent cells with Type I versus II vestibular responses (i.e., preferred direction was ipsilateral versus contralateral, respectively). *Insets*: polar plots where the vector length and angle represent each neuron’s vestibular response sensitivity and phase, respectively. Filled and open arrows represent the population-averaged vectors for Type I and II cells, respectively.

To quantify the vestibular sensitivity of each Purkinje cell in our population we fit a least-squares dynamic regression model with three kinematic terms (i.e., head-in-space position, velocity, and acceleration) to responses for movements in each direction (**Figure 1—figure supplement 1**, see Methods). We found that the preferred movement direction (i.e., the direction that resulted in an excitatory response, or in the greater excitatory response in the case of v-shaped neurons) could be either ipsilateral or contralateral for a given Purkinje cell. Neurons with preferred responses for ipsilaterally (for example, **Fig. 1A**, right panel (n = 32) or contralaterally (**Fig. 1A,** left and middle panels n = 41) directed rotations were accordingly classified as Type I or Type II, respectively. Our analysis further revealed that the response dynamics varied considerably across neurons, with 43% neurons demonstrating responses that were relatively in-phase (± 15°) with head velocity (1.8 ± 7.7°), while others demonstrated marked response leads (32%, 57± 31) or lags (25%, −47.5 ± 22°). **Figures 1B,C** illustrate the vestibular response vectors for our populations of neurons computed in their preferred and non-preferred directions, respectively. The vector represents the gain (length) and phase (angle) of the neural responses to each stimulus computed at 1Hz (see Methods). The large arrows represent average neuronal responses to vestibular stimuli for Type I (S_vest._ = 0.42 ± 0.37 (sp/s)/(°/s), Phase_vest._= 6 ± 31) and Type II neurons (S_vest._ = 0.31 ± 0.34 (sp/s)/(°/s), Phase_vest._= 172 ± 42°), respectively.

We next addressed whether the Purkinje cells that responded to vestibular stimulation also responded to the activation of neck proprioceptors. To test this, we recorded neuronal activities during a paradigm in which neck proprioceptive stimulation was delivered in isolation using the same motion profile as that used above to assess neuronal vestibular sensitivities. **Figure 2A** illustrates the responses recorded from the same three example neurons shown in **Figure 1A**, while we rotated the monkey’s body beneath its earth-stationary head. We quantified the neck proprioceptive sensitivity of each Purkinje cell using least-squares dynamic regression (see Methods) and found that most neurons ~75% (n = 54) were sensitive to passive proprioceptive stimulation (**Fig. 2B&C**, filled bars; bimodal neurons), whereas the remaining ~25% (n = 19) were insensitive (**Fig. 2 Fig. 2B&C,** open bars; unimodal neurons). Furthermore, similar to our findings above regarding vestibular sensitivities, the dynamics of responses to proprioceptive stimulation varied considerably across Purkinje cells (**Figure 2—figure supplement 1).** Further, Purkinje cells showed heterogeneity in their simple spike responses to vestibular versus proprioceptive stimulation **(Figure 2—figure supplement 2).** Insets in **Figures 2B,C** illustrate the response vectors for proprioceptive stimulation across our populations of neurons in the preferred and non-preferred directions, respectively. As in **Figures 1B,C** above, the vector was computed based on the gain (length) and phase (angle) of the neural responses to each stimulus at 1 Hz (see Methods) and large arrows represent average neuronal responses measured in response to proprioceptive stimuli for neurons with Type I (S_prop._ = 0.12 ± 0.44 (sp/s)/(°/s), Phase_prop._= 159 ± 29°) versus Type II (S_prop._ = 0.13 ± 0.46 (sp/s)/(°/s), Phase_prop._= −27 ± 20°) responses.

**Figure 2.**
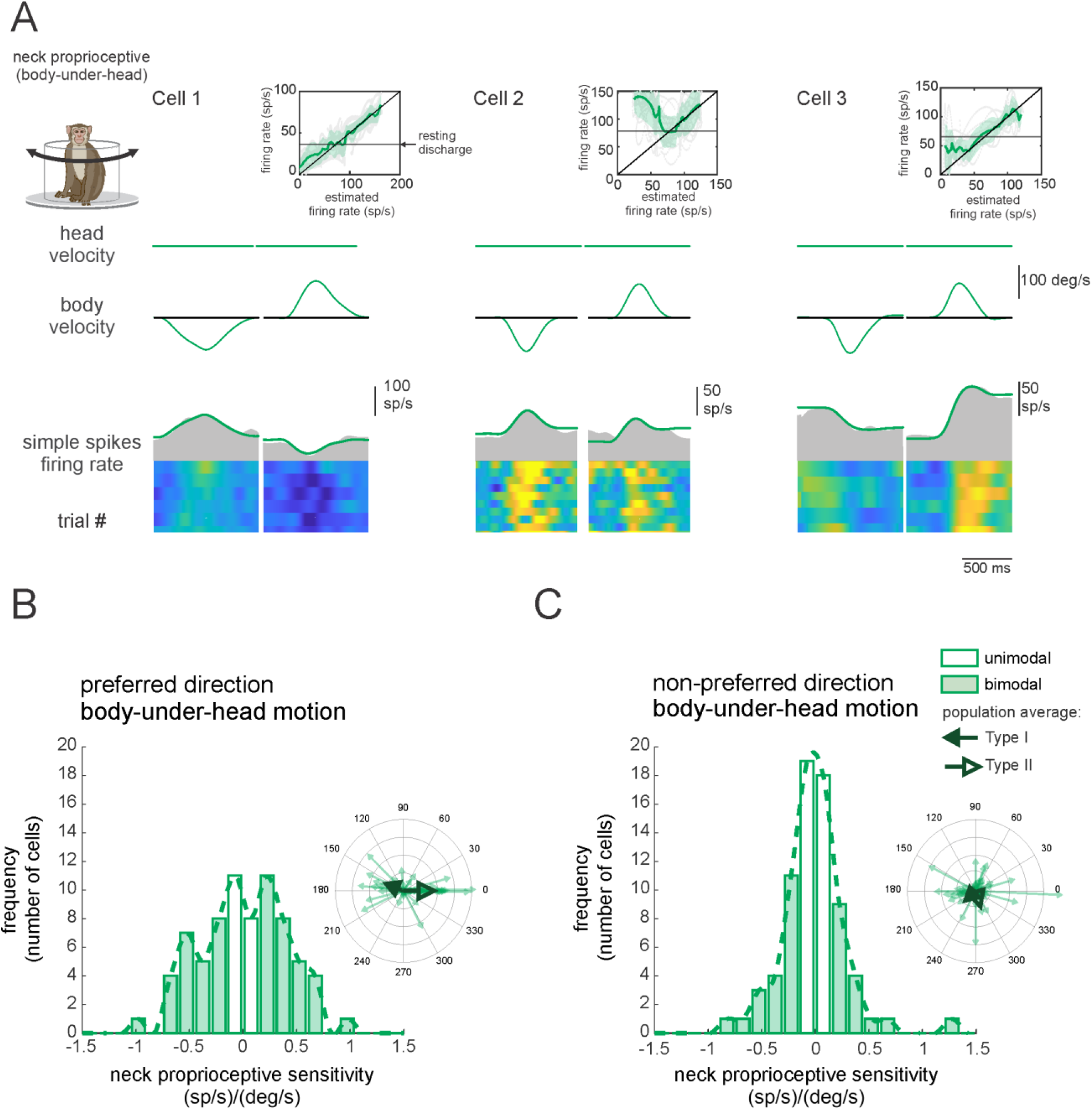
Purkinje cell simple spike responses to neck proprioceptive stimulation. **(A)** Proprioceptive stimulation was generated by applying body under head rotation about the vertical axis while holding the head earth fixed (i.e., proprioceptive stimulation condition). The resulting neural responses are illustrated for the same three example Purkinje cells shown above in **Fig. 1**. The top two rows illustrate rotational head versus body velocities. The bottom row shows the resultant simple spike firing rate (gray shaded regions) with the linear estimation of the firing rate based on body motion superimposed (green traces). Insets: the relationship between simple spike firing rate (phase-corrected) and angular body-in-space velocity. **(B, C)** Distribution of proprioceptive sensitivities for the preferred **(B)** and non-preferred **(C)** directions of body movement. Filled versus open bars represent neurons that were sensitive versus insensitive to neck proprioceptive stimulation (i.e., bimodal versus unimodal cells, respectively). *Insets*: polar plots where the vector length and angle represent each neuron’s proprioceptive response sensitivity and phase, respectively. Filled versus open arrows represent the population-averaged vectors for neurons with Type I versus II vestibular responses (i.e., **Fig. 1**), respectively.

### Purkinje cell responses to simultaneous proprioceptive and vestibular stimulation

So far, we have shown that most Purkinje cells in our population were sensitive to neck proprioceptive as well as vestibular stimulation, and that we categorized these neurons as ‘bimodal’ versus neurons that were only responsive to vestibular stimulation as ‘unimodal’. The vestibular sensitivities of bimodal Purkinje cells were comparable to those of their unimodal counterparts (*p* = 0.17). We further found that the neck sensitivities of bimodal neurons were most often (67%) antagonistic relative to their vestibular sensitivities in the *preferred* direction. This can be seen in **Figures 3A,B**, where the average vectors representing the neuronal response to the vestibular and proprioceptive stimulation point in opposite directions (**Figs. 3A,B**, thick blue versus green arrows, respectively). In contrast, relative to vestibular sensitivities in *non-preferred* direction bimodal cells were as likely to have antagonistic as agonistic responses to neck proprioceptive stimulation (**Figure 3—figure supplement 1**).

**Figure 3.**
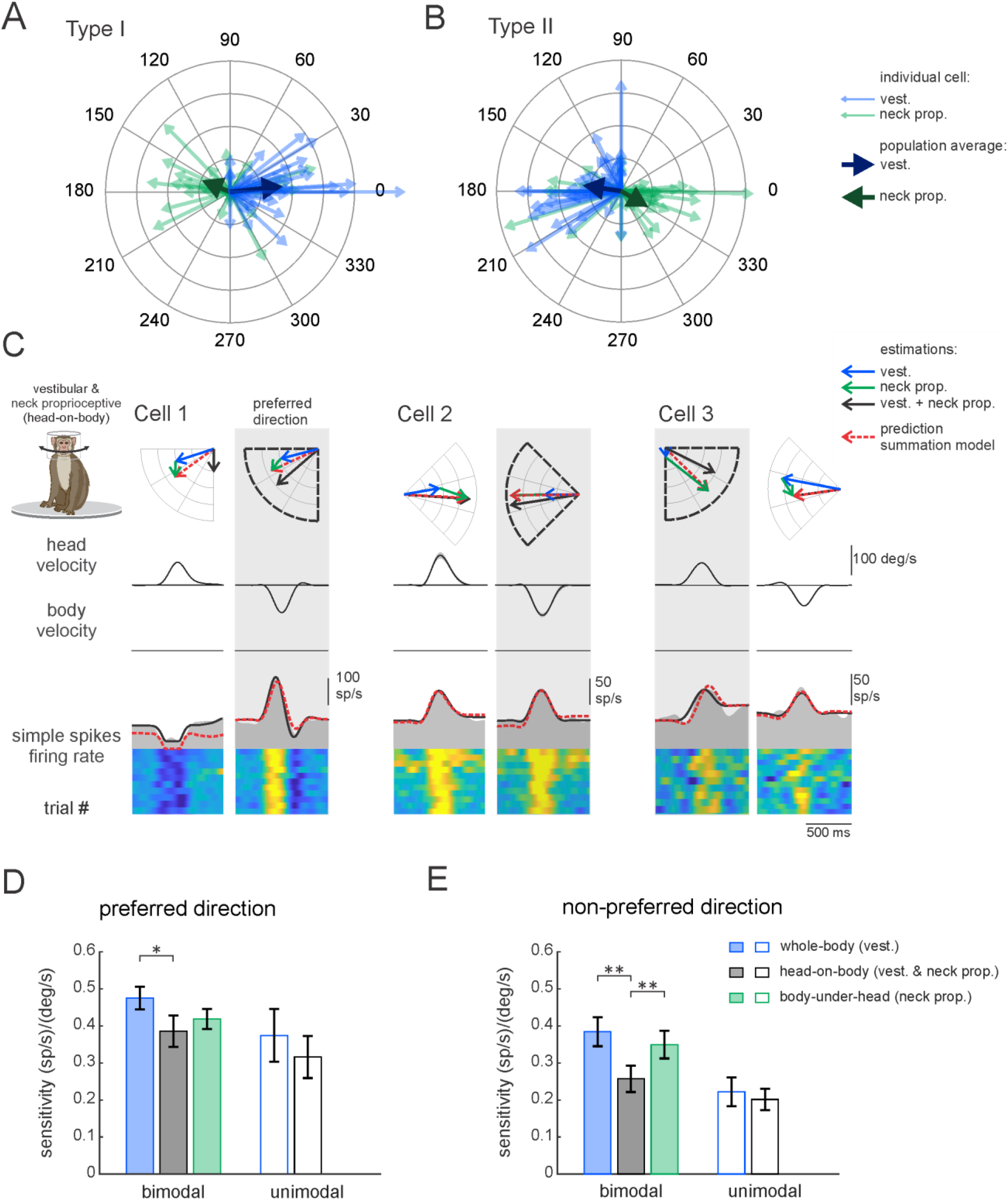
Purkinje cells simple spike responses to combined vestibular– proprioceptive stimulation. (**A, B)** Polar plots illustrating the vestibular (blue) and neck proprioceptive (green) neuronal response sensitivities of Type I **(A)** and Type II **(B)** Purkinje cells for preferred direction of vestibular stimulation and complementary direction proprioceptive stimulation (i.e., body-under-head motion). Bold blue and green arrows represent the mean population vectors, respectively. **(C)** Combined vestibular-proprioceptive stimulation was generated by applying passive head on body rotations about the vertical axis. The resulting neural responses are shown for the same three example Purkinje cells shown above in Figs. 1,2. The top two rows illustrate rotational head versus body velocity. The bottom row shows the resultant simple spike firing rate (gray shaded regions). The linear estimation of firing rate based on head motion (solid black traces) and the firing rate prediction based on the linear summation of neck proprioceptive and vestibular sensitivities (dashed red traces) are both superimposed. Each neuron’s preferred motion direction for vestibular stimulation is indicated by the gray column. Polar plots (***top***) represent the sensitivity and phase of each neuron’s response to vestibular, proprioceptive, and combined stimulation as well as the response predicted by the summation model. **(D, E)** Bar plots comparing the sensitivities of bimodal and unimodal Purkinje cells to vestibular, proprioceptive, and combined stimulation in the *preferred* **(D)** and *non-preferred* **(E)** motion directions, as defined by each neuron’s responses to vestibular stimulation. The response sensitivities of Type I and II neurons are reported as positive values relative to ipsilaterally and contralaterally directed head movements, respectively, to facilitate comparison across all Purkinje cells.

During everyday activities, we move our head relative to our body, and thus simultaneously activate both vestibular sensors and neck proprioceptors. To directly establish how vestibular and neck proprioceptive information is integrated in the anterior vermis, we next recorded the responses of the same Purkinje cell populations while the head was rotated so that it moved relative to an earth-stationary body (i.e., head-on-body rotation condition). The responses of the same three example neurons above in **Figures 1,2** are shown in **Figure 3C**. The head motion-based linear estimation of firing rate (solid black traces, see Methods) is plotted on the firing rate for each cell. A prediction of firing rate based on the linear summation of each neuron’s sensitivity to neck proprioceptive and vestibular stimulations when applied alone (dashed red traces) is superimposed for comparison. The example neurons were typical in that each neuron’s modulation for combined stimulation in the *preferred direction* (i.e., grey columns) was well predicted by the linear summation of the neuron’s vestibular and proprioception sensitivities. The polar plots show the sum (dashed red arrow) of the example neuron’s response vectors for the vestibular and proprioceptive stimulation (blue and green arrows) when applied in isolation versus the response vector for combined stimulation (black arrow). Correspondingly, there was good alignment between the vector length and direction computed for the firing rate estimate and prediction for these three example neurons for the head-on-body motion in the *preferred direction.*

**Figure 3D,E** summarizes the population data for unimodal and bimodal Purkinje cells in each of the three stimulation conditions. Average response sensitivities are shown for *preferred* (**Fig. 3D**) and *non-preferred* (**Fig. 3E**) direction motion. Note that since there were no significant differences between the response of Type I and II cells (other than their preferred direction) we reported the responses of both groups together, by accounting for the difference in the direction of the modulation of Type II cells. We first hypothesized that the vestibular sensitivity of unimodal neurons should remain constant across conditions regardless of whether the neck proprioceptors were stimulated. Consistent with this proposal, we found that unimodal cell response sensitivities (open bars) were comparable during the vestibular-only and combined stimulation conditions (**Figs. 3D, E**, p >0.22). Likewise, response phases of unimodal neurons were comparable for both conditions (*preferred*: 11±5 versus 11±14; p=0.21; vs. *non-preferred*: 27±8 versus 14±18; p=0.77). In contrast, we hypothesized that since the vestibular and proprioceptive sensitivities of bimodal Purkinje cells were generally antagonist (i.e., **Fig. 3A, Figure 3—figure supplement 1**), the oppositely modulated inputs from neck proprioceptors should effectively suppress the vestibular driven responses during the combined conditions. Indeed, consistent with this prediction, the sensitivities of bimodal Purkinje cells were reduced during the combined stimulation condition relative to the vestibular-only condition **(Figs. 3D,E**, filled bars, p <0.027).

Above, we showed that the responses of our three example neurons in the combined stimulation were well predicted by the linear summation of the neuron’s vestibular and proprioception sensitivities, particularly for stimulation in the *preferred direction*. We next explicitly addressed whether this simple linear model of vestibular-proprioceptive integration could reliably predict neuronal responses in the combined condition across our population of Purkinje cells. To do this, we compared on a neuron-by-neuron basis and estimated and predicted head-on-body rotation sensitivities (**Fig. 4A**) and phases (**Fig. 4B**) for all the Purkinje cells in our sample. Overall, estimated and predicted sensitivities and phases were comparable (R^2^ = 0.77 and 0.75 for sensitivity and phase, respectively, p<0.001). The similarity between values is shown by the slope of the line fits to the data which were not different from 1 (p = 0.42 and 0.83 for gain and phase, respectively). Thus, the summation model provided a good estimate of the gain of the preferred direction responses during combined stimulation. Similar results were obtained in our analysis of the non-preferred direction responses (**Figure 4—figure supplement 1**).

**Figure 4.**
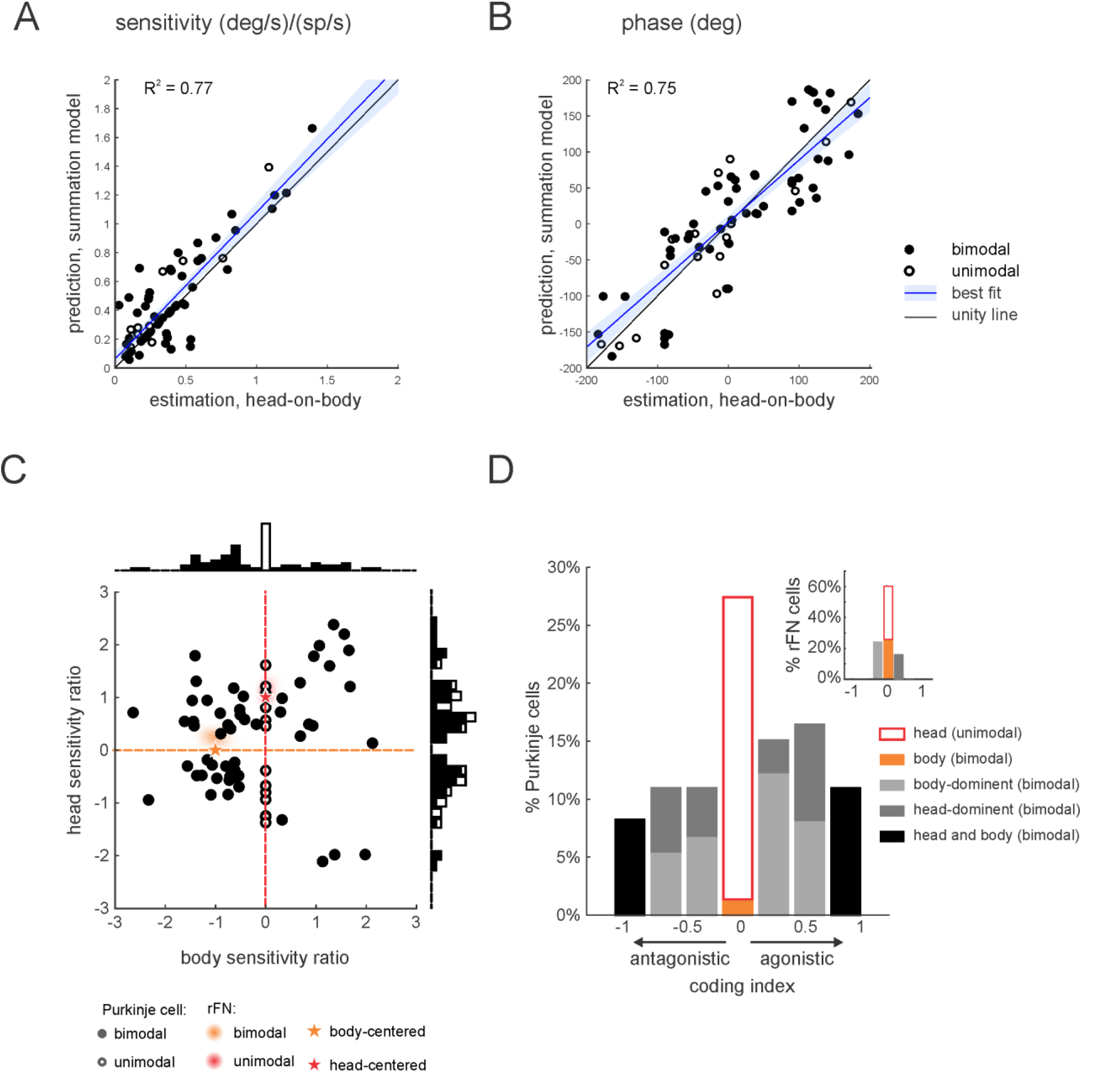
Purkinje cell simple spike responses to combined stimulation are well predicted by the linear summation of a given neuron’s responses to vestibular and proprioceptive stimulation when applied alone. **(A,B)** Comparison of estimated and predicted sensitivities (**A**) and phases (**B**) of Purkinje cell responses to head-on-body rotations in the preferred movement direction. The linear summation of a given neuron’s vestibular and neck proprioceptive sensitivities well predicted both measures in the combined condition. Blue lines and shading denote the mean ± 95% CI of linear fit. **(C)** Scatter plot of the relationship between the head sensitivity ratio (S_vest._+prop. / S_vest._) and body sensitivity ratio (S_prop._ / S_vest._). Histograms (top and right) illustrate the distributions of body and head sensitivity ratios, respectively. Orange versus red stars indicate ideal neurons encoding body versus head movement in space, respectively. For comparison, the red and orange shaded areas representing the distribution of values estimated for unimodal and bimodal rFN neurons (Brooks et al. 2009) are superimposed. **(D)** Distribution of coding indexes (see Methods). Positive and negative values correspond to agonistic and antagonistic responses to head vs. body encoding, respectively. Inset: the distribution of coding indices estimated for rFN neurons (Brooks et al. 2009) is shown for comparison.

As stated above, the generation of vestibulospinal reflexes requires central pathways to explicitly transform vestibular information from a head-centered to a body-centered reference frame during self-motion. To better understand the coding by our population of neurons, we first computed a “head sensitivity” and a “body sensitivity” ratio for each neuron (**Fig. 4C**, see Methods). Two theoretical neurons, one that selectively encodes head-in-space and the other that selectively encodes body movement are indicated by red and orange stars respectively. A neuron selectively encoding head-in-space motion (red star) would display comparable responses to whole-body and head-on-body rotations (head sensitivity ratio= 1), while not responding to body-under-head rotations (body sensitivity ratio= 0). On the other hand, a neuron selectively encoding body motion would display a comparable response to whole-body and body-under-head rotations (orange star, body sensitivity ratio= 1), while not responding to head-on-body rotations (head sensitivity ratio= 0). In contrast, comparison of these ratios across each of the Purkinje cells in our neuronal population revealed considerable heterogeneity in the relationship between these two measures relative to these theoretical neurons. As reviewed above, anterior vermis Purkinje cells target neurons in the deep cerebellar nuclei (i.e., rFN) (Batton et al., 1977; Yamada and Noda 1987). Thus, for comparison, we computed these ratios for rFN neurons that had been studied during the same conditions (Brooks and Cullen, 2009). The red and orange shaded areas represent the distribution of our sensitivity ratios for the unimodal and bimodal rFN neuron populations reported in this prior study. Notably, in contrast to the Purkinje cells of our present study, the relationship between the rFN unimodal and bimodal neuron sensitivity ratios are well-aligned with that of our theoretical neurons that selectively encoded head and body movement, respectively.

To next evaluate the transformation of vestibular information from a head-centered to body-centered reference frame during self-motion for each Purkinje cell we computed a coding index (see Methods). Specifically, this index compared each neuron’s sensitivity when only the head moved relative to space (i.e., head-on-body rotation, **Fig. 3**) versus when only the body moved relative to space (i.e., body-under-head rotation, **Fig. 2**), to the combined stimulation condition. The results from the analysis of our Purkinje cell population are shown in **Figure 4D**. Indeed, only a minority of neurons were designated as primarily head (26%) or body (2%) encoding (light and dark orange bars, respectively). A complementary analysis of *non-preferred* direction responses (relative to vestibular stimulation) revealed similar results (**Fig. 4—figure supplement 1**). Again, for comparison, the corresponding distribution of coding indices estimated for rFN neurons from Brooks et al. 2009 is shown for comparison (**Figure 4D,** top right inset). In contrast to Purkinje cell’s the majority of neurons were designated as primarily head (34%) or body (26%) encoding.

### Influence of head position on Purkinje cell vestibular responses

In theoretical models of reference frame transformations, responses to the sensory inputs are generally modulated by a postural signal (e.g., the position of the head relative to the body) (Pouget and Snyder, 2000, Salinas 2001). Indeed, neurons in the deep cerebellar nuclei of primates show such tuning. Specifically, the vestibular responses of bimodal neurons in the rostral fastigial nucleus modulate as a function of head position (Brooks and Cullen 2009; Kleine et al., 2004; Shaikh et al., 2004)). This finding has been taken as support for the view that a reference frame transformation of vestibular signals from head- to body-centered occurs in the cerebellar vermis (reviewed in Cullen 2019). Thus, we next asked: How is this tuning generated? And more specifically, is it computed within the deep cerebellar nuclei or instead inherited from the Purkinje cells that target neurons in the deep cerebellar nuclei? To address these questions, we first determined whether the vestibular responses of Purkinje cells were affected by static changes in head-on-body position. We measured neuronal responses to vestibular stimulation (i.e., whole-body rotation) applied with the head positioned at 5 different orientations ranging from −30 (left) to +30° (right) relative to the body (−30, −15, 0, 15, and 30°). The example bimodal neuron was typical in that it displayed marked changes in vestibular sensitivity with changes in head head-on-body position (**Fig. 5A**). In contrast, we did not find evidence for such tuning in unimodal neurons (**Fig. 5B**).

**Figure 5.**
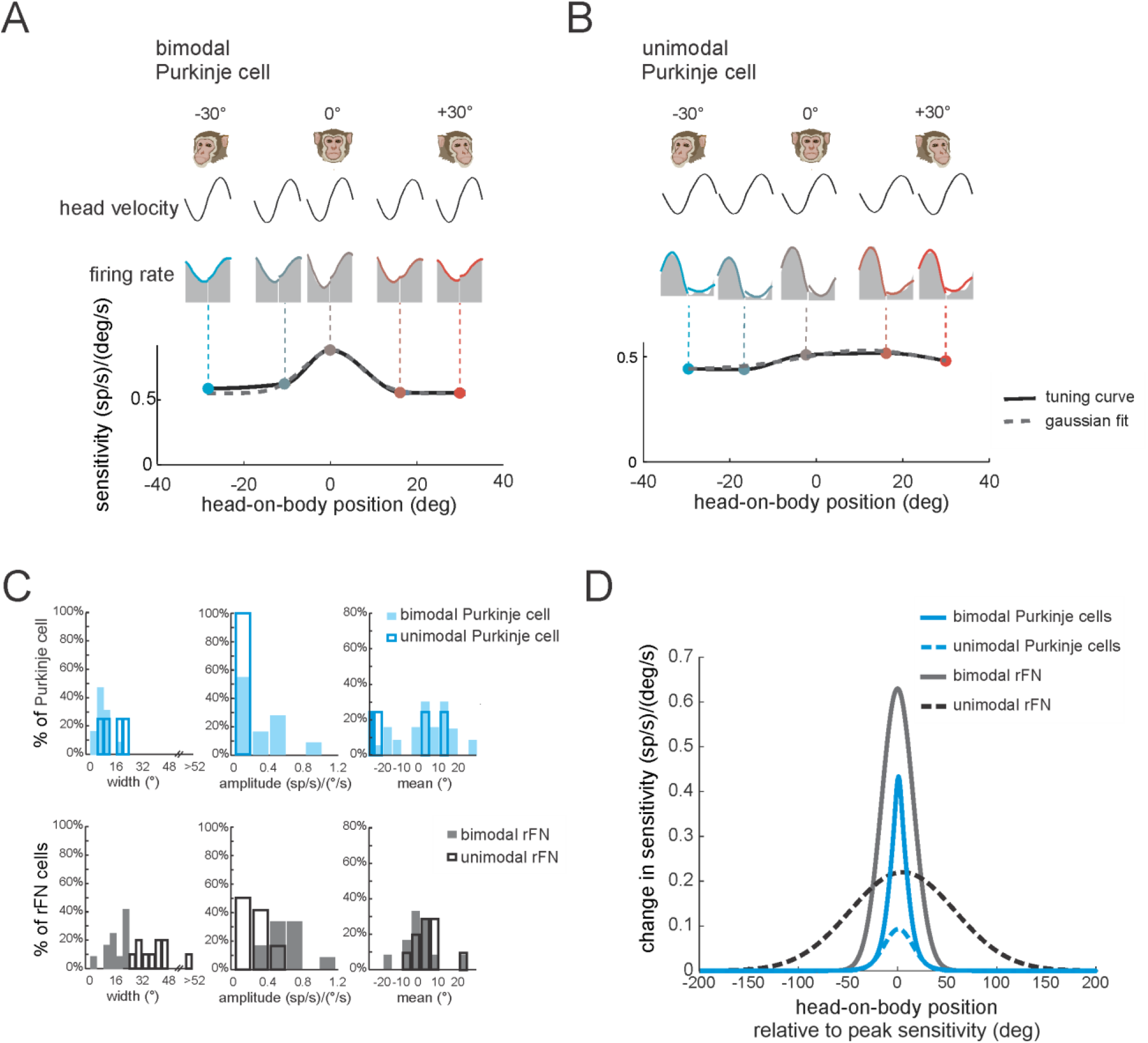
The vestibular responses of bimodal Purkinje cells show head-on-body position dependent tuning. **(A, B)** Tuning curves for the vestibular sensitivities of an example bimodal **(A)** and unimodal **(B)** Purkinje cell measured by applying whole-body rotation with the head oriented at different positions relative to the body. Note, bimodal neurons, but not unimodal neurons, show tuning as a function of head-on-body position. *Top panel:* Distributions of tuning widths (left), amplitudes (middle), and means (right) for bimodal (filled bars, N=12) and unimodal (open bars, N=5) Purkinje cells. *Bottom panel:* For comparison, the same distributions are plotted for a population of rFN neurons previously characterized using a comparable approach (Brooks and Cullen 2009). **(D)** Average tuning curves computed by aligning the peak of each individual neuron’s tuning curve. Average tuning curves are shown for bimodal and unimodal Purkinje cells (blue) for vestibular stimulation with the head oriented at different positions relative to the body. Again, for comparison, the average tuning curves of rFN neurons are superimposed (gray, data from Brooks and Cullen, 2009).

To quantify each Purkinje cell’s tuning we fit a Gaussian function to vestibular sensitivity as a function of head position (see Methods), and computed the tuning width, amplitude, and mean direction provided by the best fit to each neuron (**Fig. 5C**, top row; filled and open blue bars denote bimodal and unimodal neurons). First, bimodal neurons were more narrowly tuned than were unimodal neurons (mean tuning widths; 7.2 vs. 15°, respectively). Additionally, bimodal neurons showed stronger tuning relative to unimodal neurons (mean tuning amplitude; 0.52 vs. 0.05 (sp/s)/(deg/s), respectively. Finally, there was no difference in the mean of the tuning curve between unimodal neurons and bimodal neurons (p > 0.37). We next compared the tuning of our bimodal and unimodal Purkinje cells with that previously described for their target neurons in the rFN (Brooks and Cullen, 2009). The corresponding distributions of rFN neuron tuning width, amplitude, and mean direction are plotted in the bottom row of **Fig. 5C**. To facilitate comparison between the tuning of Purkinje and rFN cells, we aligned the peak of each individual neuron’s tuning curve with zero and averaged the resultant curves across bimodal and unimodal groups for each (**Fig. 5D**). Overall, the strength of tuning was significantly higher for bimodal rFN than Purkinje cells (**Fig. 5D** compare solid gray and black lines, ~30% reduction for Purkinje cells, p<0.001). Tuning width was also reduced for bimodal Purkinje cells (~40% reduction), while mean tuning direction was comparable for both cell groups (*p* > 0.05). Moreover, tuning was consistently stronger for bimodal than unimodal neurons in Purkinje cells as has previously been shown for rFN neurons (**Fig. 5D**, compare solid and dashed lines). The significance of these results will be addressed in the Discussion below.

### Linear combination of the Purkinje cells’ response can encode head and body motion

To summarize, our results have shown that while most vestibular-sensitive Purkinje cells in the anterior vermis integrate vestibular and neck proprioceptive signals, the transformation from head- to body-centered reference frame is not complete. Instead, single bimodal Purkinje cells generally dynamically encoded intermediate representations of self-motion that were between head and body motion. In contrast, bimodal neurons in the deep cerebellar nuclei - the primary target of these Purkinje cells (**Fig. 6A;** rostral fastigial nucleus) - dynamically encode body motion (i.e., orange shaded region, **Fig. 4C**) and also show stronger vestibular tuning as a function of head-on-body-position. Thus, taken together, our present results suggest that the transformation from a head- to body-centered representation of self-motion is achieved by integrating the activities of multiple Purkinje cells.

**Figure 6.**
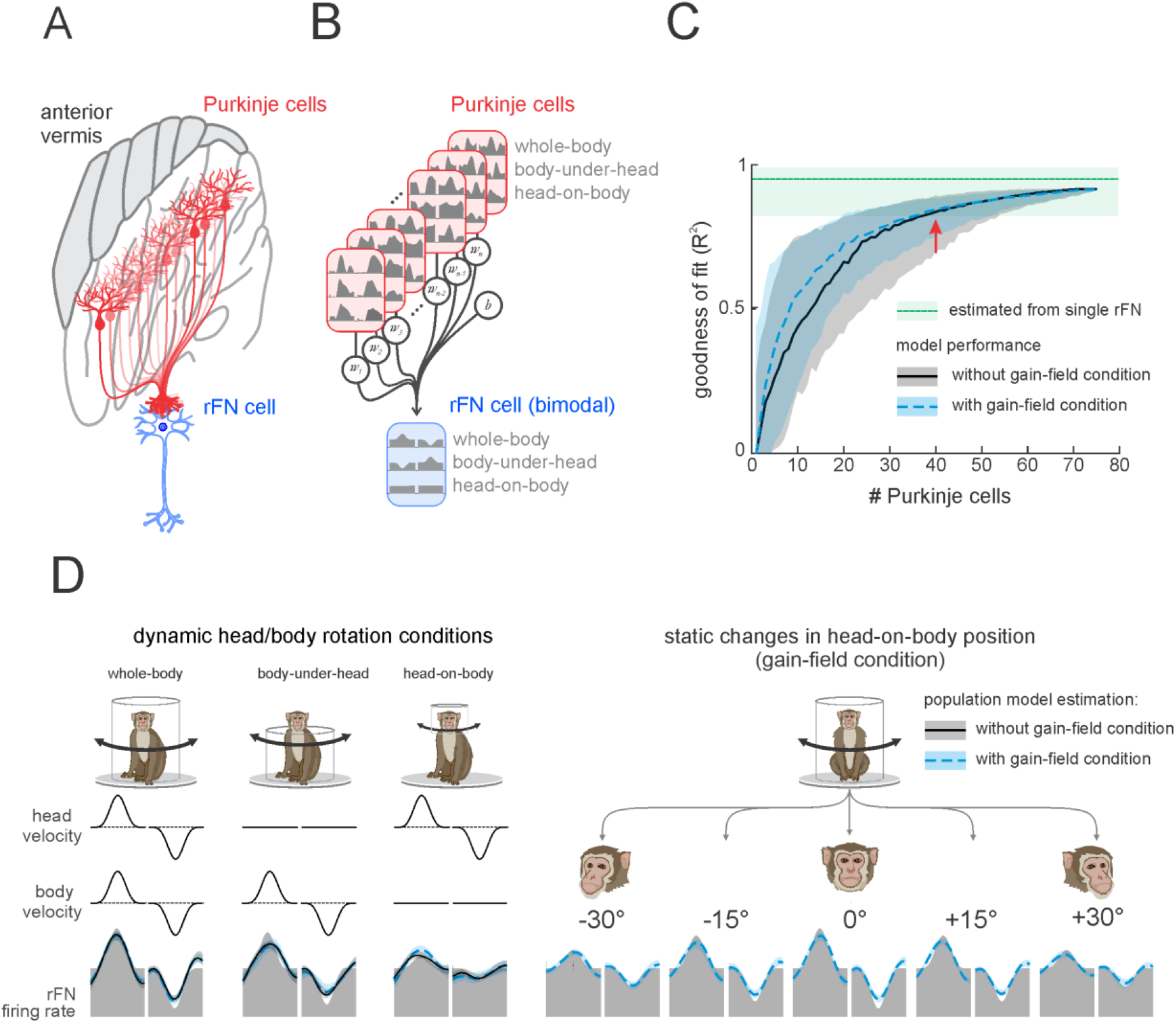
A simple linear population model of Purkinje cell integration can explain the responses of target bimodal neurons in deep cerebellar nuclei across all self-movement conditions. **(A)** Illustration of the convergence of multiple Purkinje cells onto a single neuron in the rostral fastigial nucleus (rFN), with different shades of red representing theoretical differences in the weighing of each Purkinje cell’s synapse with the target rFN neuron. **(B)** Schematic of the linear summation population model used to estimate the firing rate of a target neuron in the rFN. Each Purkinje cell’s weight was optimized to generate the best estimate of the average bimodal rFN neuron across conditions (Brooks and Cullen 2009). (**C**) Model performance as a function of the number of Purkinje cells. Black curve corresponds to model fit to the simple spike firing rates of all 75 Purkinje cells recorded during three passive conditions (i.e., whole-body, body-under-head, and head-on-body movements. Blue curve corresponds to the model fit to simple spike firing rates of all 75 Purkinje cells during our three passive conditions as well as simulated responses of these cells recorded in the gain-field (static head-on-body position; **Fig. 5**). The variability estimated from a population of rFN bimodal neurons previously described by Brook and Cullen (2009) is represented by the green shaded band. **(D)** Estimated model firing rates based on a population of 40 Purkinje cells superimposed on the actual average firing rate of a bimodal rostral fastigial nucleus (rFN) neuron (grey shaded region). Solid black lines versus dashed blue lines illustrate firing rate estimations from models that included (i) the three dynamic head/body rotation conditions (left) versus (ii) the three dynamic conditions as well as the gain-field condition (right).

Accordingly, we next tested this hypothesis. Specifically, to quantify the actual number of Purkinje cells necessary to explain the responses of bimodal rFN neurons, we determined whether a simple linear model optimizing the weights of the activities of multiple Purkinje cells (see Methods) could generate bimodal rFN neural responses across conditions (**Fig. 6B**). As expected, combining the activities of more Purkinje cells (i.e., increasing population size) led to an increase in the goodness of fit (**Fig. 6C**). Data sets used for the modeling included Purkinje cell responses during (i) our three dynamic conditions (i.e., whole-body, body-under-head, head-on-body rotations) alone (black curve), and (ii) these same three dynamic conditions as well as the static changes in head-on-body position (i.e., **Fig. 5**) condition (dashed blue curve). Importantly, in both cases we found that the weighted activities of ~40 neurons generated responses that well approximated those previously reported for bimodal rFN neurons (**Fig. 6C,** red arrow); the confidence intervals our model estimates that of the rFN neural responses completely overlapped for a population of ~40 neurons. Thus, a population of 40 Purkinje cells could explain the dynamic representation of body motion across conditions (**Fig. 6D**), as well as robust encoding of vestibular stimuli as a function of static head position observed in bimodal rFN neurons (**Fig. 6D,** right panel). For completeness, we also used the same approach to quantify the number of Purkinje cells necessary to explain the responses of unimodal rFN neurons and obtained comparable results (**Fig. 6—figure supplement 1**). Interestingly, our finding that a population of 40 Purkinje cells is again required to explain the responses of bimodal and unimodal rFN neurons matches the value established independently from anatomical studies of Purkinje cell - deep cerebellar nucleus neuron projection ratio. We further consider this point below in the Discussion.

## Discussion

### Summary of results

Here we recorded the simple spike activity of Purkinje cells of the anterior vermis during passive vestibular (i.e., whole-body rotation), neck proprioceptive (i.e., body-under-head rotation), and a combination of vestibular and neck proprioceptive stimulation (i.e., head-on-body rotation). First, we found that most Purkinje cells responded to both vestibular and neck proprioceptive stimulation (i.e., bimodal neurons). Second, the linear combination of the responses to dynamic neck proprioceptive and vestibular stimulation alone provided a good estimate of each Purkinje cell’s response during combined stimulation. Third, bimodal neurons generally did not encode either the motion of the head or body in space across conditions. Instead, they dynamically encoded intermediate representations of self-motion between head and body motion. Additionally, bimodal neurons, but not unimodal neurons, showed tuning for the encoding of vestibular stimuli as a function of static head position. Finally, using a simple linear population model, we establish that combining inhibitory responses from ~40 Purkinje cells can explain the responses of target neurons in deep cerebellar nuclei across all self-movement conditions. Thus, our findings in alert monkeys provide new insight into the neural mechanisms underlying the coordinate transformation by which the cerebellum uses neck proprioceptive information to transform vestibular signals from a head- to body-centered reference frame.

### Purkinje cells have diverse temporal responses to dynamic vestibular and neck proprioceptive sensory stimulation

The integration of vestibular and neck proprioceptive-related information is required to convert head-centered vestibular signals to the body-centered reference frame required for postural control (reviewed in Cullen 2019). Our recordings in the anterior vermis of alert monkeys demonstrate that most vestibular sensitive Purkinje cells also encode neck proprioceptive-related information. As reviewed above, anterior vermis Purkinje cells project to the rostral fastigial nucleus (rFN), the most medial of the deep cerebellar nuclei (Fujita et al. 2020, Husson et al. 2014) which plays a key role in the control of posture. Two types of rFN neurons have been previously identified in alert monkeys: unimodal and bimodal neurons (Brooks and Cullen 2009). Unimodal neurons respond to vestibular stimulation during passive rotations and dynamically encode head movement. Bimodal rFN neurons respond to both vestibular and neck proprioceptive stimulation and dynamically encode body movement. Notably, because the vestibular and neck proprioceptive sensitivities of rFN bimodal neurons are both equal and complementary in sign, they sum linearly to effectively cancel each other during passive head-on-body rotations - a condition in which both sensory systems are activated but the body does not move in space. In contrast, here we found that vestibular-sensitive anterior vermis Purkinje cells were on average more sensitive to vestibular than proprioceptive stimulation and that there was considerable variability in the relative signs of responses to each modality. As a result, cancellation of these two inputs during passive head-on-body rotations was the exception rather than the rule (i.e., **Figure 3B**) with the vast majority of bimodal Purkinje cells demonstrating significant modulation in response to passively applied head-on-body rotations. Thus, unlike bimodal rFN neurons, which dynamically encode body motion, bimodal Purkinje cells dynamically encode an intermediate representation of self-motion.

### Reference frame transformations: Purkinje cell vestibular responses modulated by posture

Theoretical models of reference frame transformations commonly include a sensory input (e.g., vestibular or visual information) that is modulated by a postural signal (e.g., head-on-body position) (Pouget and Snyder 2000; Salinas 2001). The resultant modulation of the sensory signal is commonly referred to as a gain field (Andersen and Mountcastle 1983) and is thought to be mediated via nonlinear interactions between sensory responses and head/body referenced cues (Zipser and Andersen, 1988; Salinas 2001). Our present results reveal the neural substrate of such a reference frame transformation required for postural control. Notably, bimodal anterior vermis Purkinje cells displayed vestibular tuning as a function of head-on-body position during horizontal rotations. This tuning is similar but not as strong as that shown by downstream bimodal neurons in the target rFN (Brooks and Cullen 2009), and indeed some rFN neurons do encode vestibular information in a body-centered reference frame for both two dimensional (Kleine et al., 2004; Shaikh et al., 2004) and three dimensional (Green et al. 2018) self-motion. Thus, our present data establish that the modulation of vestibular information by a postural signal becomes more marked in the progression from the cerebellar cortex to the deep cerebellar nuclei. Interestingly, in the present study, such nonlinear interactions between neck position and vestibular signals were only in bimodal and not unimodal vestibular Purkinje cells. Future experiments are required to understand the implications of the dynamic coding head rather than body movements by the unimodal neurons.

Finally, it is noteworthy that our findings regarding transformation from a vestibular to head-centered reference frame in the anterior vermis of the alert primate contrast with those of Manzoni and colleagues in anesthetized decerebrate cats. Using a ‘wobble’ stimulus to dynamically tilt the head, they concluded that the direction of average response vector of Purkinje cells encoding both vestibular and proprioceptive information well corresponded to body tilt – consistent with a complete transformation from head- to body-centered reference frame (Mazoni et al., 1998; 2004). One potential explanation for this apparent difference in neural strategy is that our studies were performed in intact alert behaving animals whereas Manzoni and colleagues completed their experiments in anesthetized decerebrate preparation where modulation/gating by cortical structures is not present. Additionally, there are significant differences across species regarding how the vestibular system integrates multimodal information even at the first stage of central processing in the vestibular nuclei (reviewed in Cullen 2019). For instance, vestibular nuclei neurons in alert mice, cats, and cynomolgus monkeys commonly display vestibular–proprioceptive convergence (Medea and Cullen 2013; Cullen 2016, McCall et al., 2017). In contrast, in rhesus monkeys, vestibular nuclei neurons are only sensitive to vestibular input, and instead, proprioceptive information is integrated only at the subsequent levels of vestibular processing, most notably in the deep nuclei of the cerebellum (Roy and Cullen 2001; Brooks and Cullen 2009; Carriot et al., 2013).

### Population coding: the heterogeneous response of Purkinje cells and convergence in the rFN

Our results establish that there is considerable heterogeneity in the response dynamics of anterior vermis Purkinje cells to vestibular and/or neck proprioceptive sensory stimulation. Semicircular canal afferents and vestibular nuclei neurons provide the primary source of vestibular information to the cerebellum via mossy fiber input. However, while they encode head velocity with a phase lead, the responses of individual Purkinje cells actually more often lagged rather than led head velocity. Albus and Marrs proposed that the divergent feedforward mossy fiber projections onto a far larger number of granule cells effectively expand the dimensionality of neural space, in turn allowing better downstream decoding to linearly classify dynamic patterns of activity (Marr 1969; Albus, 1971). Indeed, recent studies have shown that mossy fibers from multiple sensory systems converge on each of more than 50 billion granule cells (Chabrol et al., 2015, Knogler et al. 2017, Lanore et al., 2021), with interneurons likely further contributing to the temporal diversity of granule cell responses (Rousseau, 2012; Kennedy et al. 2014). In turn, > 100,000 granule cells project to a single Purkinje cell via parallel fibers (Fujishima et al., 2018). Thus, together these features of the cerebellar microcircuitry are well designed to generate high-dimensional dynamic coding of information by Purkinje cells relative to their mossy fiber input across regions of the cerebellum including the anterior vermis.

In this context, the heterogeneity we observed in anterior vermis Purkinje cells responses then contrasts strikingly with the responses of their target neurons in rFN (Brooks and Cullen, 2009). Our estimation that pooling the responses of a population of ~40 Purkinje cells can explain more homogeneous responses of rFN neurons matches that established independently from anatomical studies of the Purkinje cell- deep cerebellar nucleus neuron projection ratio in rodents and cats (Person and Raman2012; Palkovits et al., 1977). Interestingly, Purkinje cells can display patterns of neuronal synchrony during active movements (Person and Raman 2012; Sarnaik and Raman 2018; Wu and Raman 2017) which could, in turn, alter the timing and modulation of target neuron responses in the deep cerebellar nuclei in a non-linear manner. Nevertheless, we found that responses could be predicted using a simple linearly weighted summation of ~40 neurons.

Finally, it is noteworthy that our study focused on the sensory responses of Purkinje cells (i.e., responses to vestibular and/or proprioceptive stimulation) during *passively* applied self-motion. Prior studies focused on the responses of Purkinje cells during *voluntary* movements have similarly concluded that they are more heterogeneous than those of their target neurons in the deep cerebellar nuclei (e.g., saccades: Thier et al., 2000; wrist control: Tomatsu et al., 2016). Interestingly, in their analysis, Tanaka et al. (2019) likewise estimated that linearly pooling the responses of ~40 Purkinje cells could account for the more homogeneous responses of target neurons in the deep cerebellar nucleus (i.e., the dentate nucleus) during voluntary wrist movements. We speculate that expanded dimensionality of the cerebellum provides a basis set for sensorimotor errors as well as plasticity at the level of Purkinje cells required to generate accurate movements (reviewed in Sohn et al. 2020) as well as ensure robust calibration over time. Overall, our current results reveal a striking transformation from heterogeneous response dynamics of cerebellar Purkinje cells to more stereotyped response dynamics of neurons in the targeted deep cerebellar nucleus. These findings provide new insights into the neural computations that ultimately ensure accurate postural control in our daily lives.

## Methods

### Experimental Model and Subject Details

Animal experimentation: All experimental protocols were approved by the Johns Hopkins University Animal Care and Use Committee and were in compliance with the guidelines of the United States National of Health. The cerebellar recordings were conducted in two male macaque monkeys (Macaca mulatta). The animals were housed on a 12hour light/dark cycle. The recording sessions were about three times a week, for approximately two hours each session. Both animals had participated in previous studies in our laboratory, but they were in good health condition and did not require any medication.

#### Method Details

##### Surgical Procedures

The two animals were prepared for chronic extracellular recording using aseptic surgical techniques described previously (Massot et al., 2012). Briefly, animals were pre-anaesthetized with ketamine hydrochloride (15 mg/kg im) and injected with buprenorphine (0.01 mg/kg im) and diazepam (1 mg/kg im) to provide analgesia and muscle relaxation, respectively. Loading doses of dexamethasone (1 mg/kg im) and cefazolin (50 mg/kg iv) were administered to minimize swelling and prevent infection, respectively. Anticholinergic glycopyrrolate (0.005 mg/kg im) was also preoperatively injected to stabilize heart rate and to reduce salivation, and then again, every 2.5–3 h during surgery. During surgery, anesthesia was maintained using isoflurane gas (0.8%– 1.5%), combined with a minimum 3 l/min (dose adjusted to effect) of 100% oxygen. Heart rate, blood pressure, respiration, and body temperature were monitored throughout the procedure. During the surgical procedure, a stainless-steel post for head immobilization and recording chambers were fastened to each animal’s skull with stainless-steel screws and dental acrylic. Craniotomy was performed within the recording chamber to allow electrode access to the cerebellar cortex. An 18-mm-diameter eye coil (three loops of Teflon-coated stainless-steel wire) was implanted in one eye behind the conjunctiva. Following surgery, we continued dexamethasone (0.5 mg/kg im; for 4 days), anafen (2 mg/kg day one, 1 mg/kg on subsequent days), and buprenorphine (0.01 mg/kg im; every 12 h for 2–5 days, depending on the animal’s pain level). In addition, cefazolin (25 mg/kg) was injected twice daily for 10 days. Animals recovered in 2 weeks before any experimenting began.

##### Data acquisition

During the experiments, the monkey sat in a primate chair secured to a turntable, and its head was centered in a coil system (CNC Engineering). Extracellular single-unit activity was recorded using enamel-insulated tungsten microelectrodes (Frederick-Haer). The location of the anterior vermis of the cerebellar cortex was determined relative to the abducens nucleus identified based on stereotypical neuronal responses during eye movements. The Purkinje cells were identified by their characteristic complex spike activity. The angular velocity of the turntable was measured using a gyroscope sensor (Watson Industries, Eau Claire, Wisconsin). Monkeys’ gaze and head angular positions were measured using the magnetic search coil technique. The neck torque produced by the monkey against its head restraint was measured using a reaction torque transducer (QWFK-8M; Honeywell, Canton, MA). All analog behavioral signals were low-pass filtered with a 125Hz cut-off frequency and acquired at 1 kHz. The neural activity was recorded at 30kHz using a data acquisition system (Blackrock Microsystems). Action potentials from the neural recording were sorted using a custom Matlab GUI (MathWorks), which provides threshold, clustering, and manual selection/removal methods.

##### Head and Body motion paradigms

Two monkeys were trained to follow a target projected onto a cylindrical screen located 60 cm away from the monkey’s head. Eye motion sensitivities to saccades and ocular fixation were characterized by having the head-restrained monkey attend to a target that stepped between horizontal positions over a range of ±30°. Neuronal responses were also tested during smooth pursuit eye movements tracking sinusoidal target motion (0.5 Hz, 40°/s peak velocity). The sensitivities of Purkinje cell simple spikes responses (n = 56) to passive head velocity were tested by applying whole-body rotations about an earth-vertical axis in the dark with velocity trajectories mimicking voluntary head movements generated to orient to targets located at ±30° left/right relative to the center (i.e., “active-like”). After a neuron’s response to the eye and head motion had been characterized, we next investigated its sensitivity to neck proprioceptive inputs by rotating the monkey’s body with this same active-like trajectory while its head was held stationary relative to space (body-under-head rotation). The third paradigm, termed “head-on-body rotation,” in which the monkey’s head was rotated relative to their stationary body was used to assess neuronal responses to the stimulation of both proprioceptive and vestibular sensors. The “head-on-body rotation” was also performed in an active-like trajectory. Finally, in a subset of neurons, testing was also done with the head initially oriented at different positions relative to the body to access whether static neck position influenced vestibular-induced modulation during whole-body sinusoidal rotation (1Hz, ± 40°/s).

##### Data analysis

###### Analysis of neuronal discharge dynamics

Data were imported into the Matlab (The MathWorks) programming environment for analysis, filtering, and processing as previously described (Dale and Cullen, 2017). Neuronal firing rate was computed by filtering spike trains with a Kaiser window at twice the frequency range of the stimulus (Cherif et al. 2008). We first verified that each neuron neither paused nor burst during saccades and was unresponsive to changes in eye position during fixation. We then used a least-squares regression analysis to describe each Purkinje cell simple spike’s response to whole-body and body-under-head rotations:

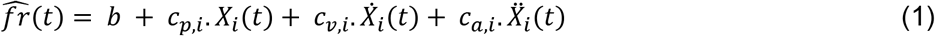

where 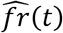 is the estimated firing rate, *b* is a bias term, *c*_*p*,*i*_, *c*_*v*,*i*_, and *c*_*a*,*i*_are coefficients representing the position, velocity, and acceleration sensitivities respectively to head (*i* = 1) or body motion (*i* = 2), and *X*_*i*_, 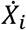 and 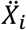 are head (*i* = 1) or body (*i* = 2) position, velocity and acceleration (during whole-body and body-under-head rotations), respectively. For each model coefficient in the analysis, we computed 95% confidence intervals using a nonparametric bootstrap approach (Carpenter and Bithell, 2000). All non-significant coefficients were set to zero. We then used coefficients to estimate the sensitivity and phase of the response using the following equations:

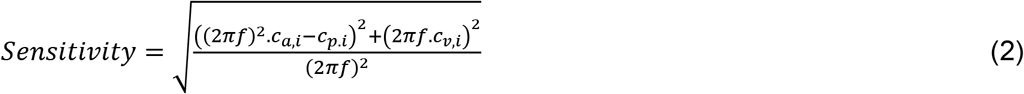

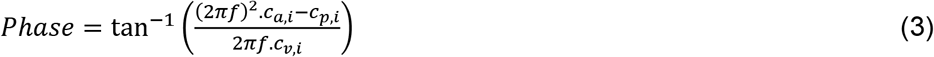

For which *f* = 1*Hz* to match the duration of half-cycle of movements (500ms). The sensitivity of the Purkinje cells to the neck proprioceptive stimulation (during body-under-head rotations) was used to categorize the cells into unimodal (zero sensitivity) and bimodal (non-zero sensitivity).

We used a similar approach to estimate sensitivities to passive head-on-body movements. Since in this condition, it is not possible to dissociate neck proprioceptive and vestibular sensitivities, we estimated them as a single coefficient. Estimated sensitivities were compared to those predicted from the linear summation of the vestibular and proprioceptive sensitivities estimated for the same neuron during passive whole-body and body-under-head rotations (termed summation model), respectively. To quantify the ability of the linear regression analysis to model neuronal discharges, the variance-accounted-for (VAF) for each regression equation was determined as previously described (Cullen et al., 1996). Values are expressed as mean ± SD and paired-sample Student’s t-tests were used to assess differences between conditions.

Due to nonlinearities observed in Purkinje cell responses, least-squares regression analysis was performed separately on firing rates recorded during ipsilateral versus contralateral rotations in each condition. We used the similarity/difference in a given Purkinje cell’s estimated sensitivity to ipsilateral versus contralateral stimulation to group it into one of four categories: (i) *linear,* which showed comparable excitatory and inhibitory sensitivities to oppositely directed head movements 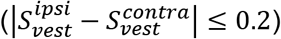, (ii) *V-shape*, which showed bidirectional excitatory responses 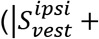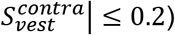 (iii) *rectifying,* which showed only excitatory responses for one movement direction and no response in the other 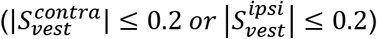, and (iv) *others*, which did not meet any of the above criteria. A given cell’s preferred direction for vestibular stimulation was characterized as the whole-body rotation direction generating the larger increase in the firing rate.

###### Quantifying head versus body encoding

We computed a ‘head sensitivity ratio’ and ‘body sensitivity ratio’ for each Purkinje cell. These ratios were defined as the neuron’s (i) sensitivity to head-on-body rotation/sensitivity to whole-body rotation, and (ii) sensitivity to body-under-head rotation/sensitivity to whole-body rotation, respectively. Further to quantify the relative encoding of head versus body motion by a given cell, we computed a ‘coding index’, which was defined as the ratio (smaller value)/(larger value) of these two ratios.

###### Quantification of head position on Purkinje cell vestibular sensitivity

The tuning curves for different head-on-body positions were fit with Gaussian curves with the following equation:

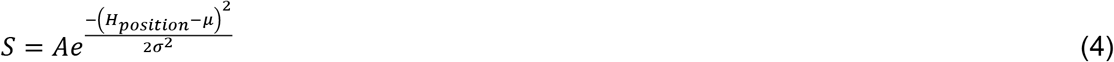

Where *μ* represents the mean, *σ* is a measure of the width, and *A* is the amplitude from the peak to the base of the Gaussian curve (as described previously, Brooks and Cullen, 2013).

###### Population modeling of Purkinje cells

To determine whether integrating the activities of multiple Purkinje cells could explain the response of their target neurons in the rFN, we used the linear model below:

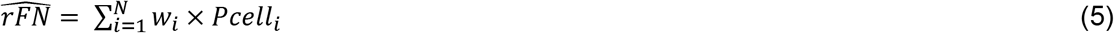

where 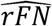 is a reconstructed firing rate response of an rFN neuron. The *w*_*i*_ corresponds to weights of connection from Purkinje cells to an rFN neuron, which all considered non-positive to reflect inhibitory synapses from Purkinje cells to rFN neurons. *Pcell*_*i*_ are observed firing rate of simple spikes *N* Purkinje cells, where *N* is a number between 1 and the total number of Purkinje cells in the dataset. For each *N* we used a bootstrapping approach to find the 95% confidence intervals of the goodness of fit (*R*^2^) as well as the model predictions.

## Acknowledgments

This research was supported by grants R01-DC002390 and R01-DC018061 from the National Institutes of Health (KEC). We would like to thank Dale Roberts for his technical support and Jessica Brooks and members of the Cullen lab for helpful discussions.

## Supplementary figure captions

**Figure 1 - figure supplement 1.**
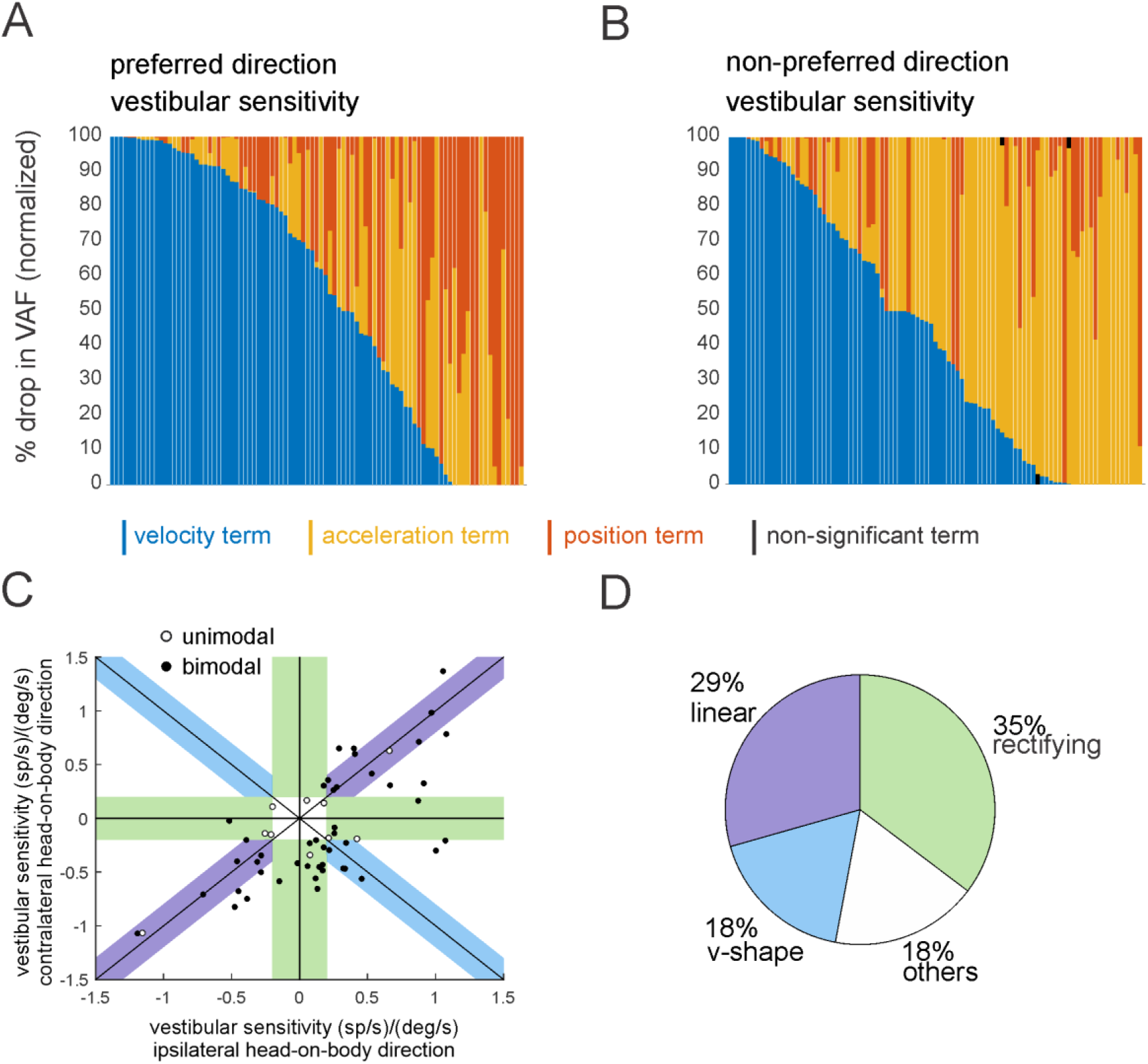
Purkinje cells show heterogeneity in their simple spike responses to vestibular stimulation. **(A, B)** The contribution of each kinematic term (i.e., position, velocity, acceleration) in estimating the firing rate for preferred **(A)** and non-preferred **(B)** direction of whole-body movement, computed as the % drop in total VAF when removed from the full model. **(C)** The response of the unimodal (white circles) and bimodal (black circles) are groups as (i) *linear* which showed excitatory versus inhibitory responses for oppositely directed head movements 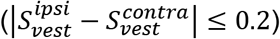, (ii) *V-shape* with bidirectional excitatory responses 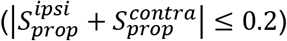, (iii) *rectifying* that generated excitatory responses for one movement 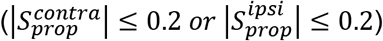, and (iv) *others* that did not meet any of the mentioned criteria. **(D)** The pie chart illustrates the percentage of each category within the Purkinje cells.

**Figure 2 - figure supplement 1.**
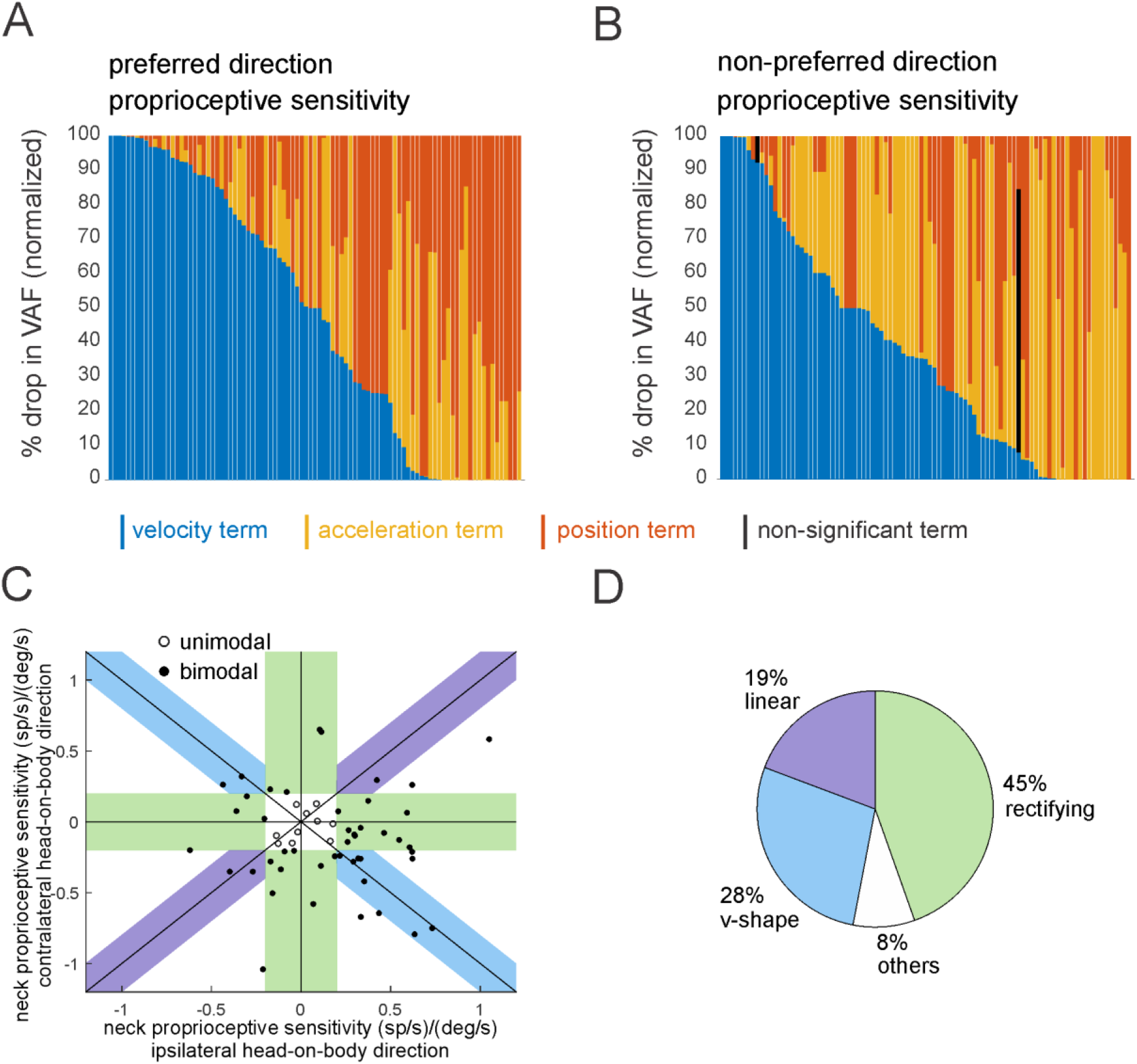
Purkinje cells show heterogeneity in their simple spike responses to proprioceptive stimulation. **(A, B)** The contribution of each kinematic term (i.e., position, velocity, acceleration) in estimating the firing rate for preferred **(A)** and non-preferred **(B)** direction of body movement (i.e., body-under-head), computed as the % drop in total VAF when removed from the full model. **(C)** The response of the unimodal (white circles) and bimodal (black circles) are groups as (i) *linear* which showed excitatory versus inhibitory responses for oppositely directed body movements 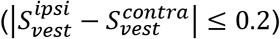, (ii) *V-shape* with bidirectional excitatory responses 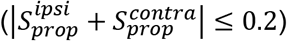, (iii) *rectifying* that generated excitatory responses for one movement 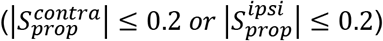, and (iv) *others* that did not meet any of the mentioned criteria. **(D)** The pie chart illustrates the percentage of each category within the Purkinje cells.

**Figure 2 - figure supplement 2.**
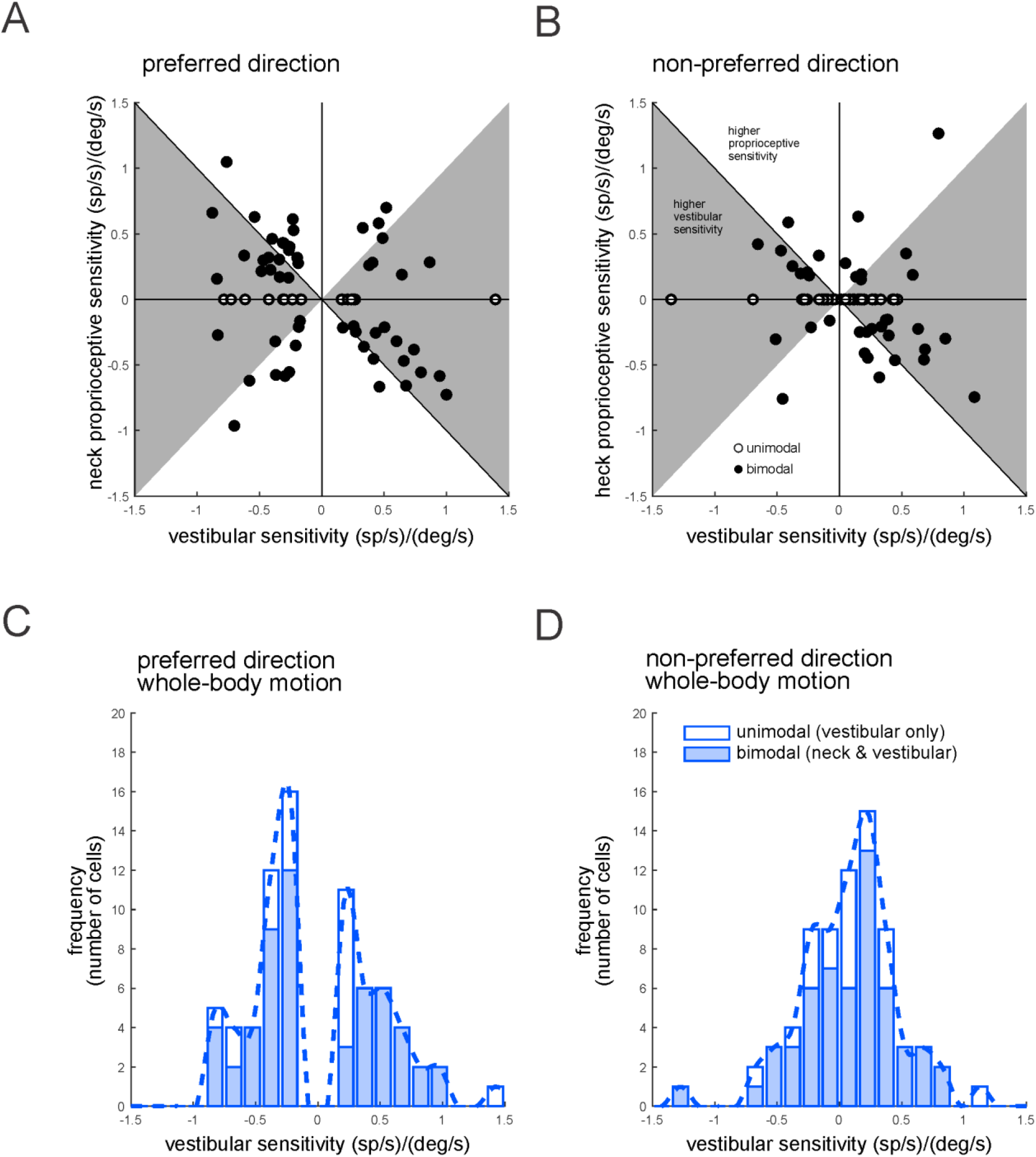
Purkinje cells show heterogeneity in their simple spike responses to vestibular versus proprioceptive stimulation. **(A, B)** Scatter plots comparing the sensitivity to vestibular and neck proprioceptive stimulation for unimodal (white circles) and bimodal (black circles) Purkinje cells for preferred **(A)** and non-preferred **(B)** direction of movement. The Grey shaded area corresponds to the Purkinje cells with larger sensitivity to vestibular compared to neck proprioceptive sensitivity. **(C, D)** The distribution of the neuronal sensitivity of the Purkinje cells simple spikes response to the vestibular stimulation for the preferred **(C)** and non-preferred (**D)** direction of movement. Open bars correspond to the cells that did not show a significant response to the neck proprioceptive stimulation (i.e., unimodal cells).

**Figure 3 - figure supplement 1.**
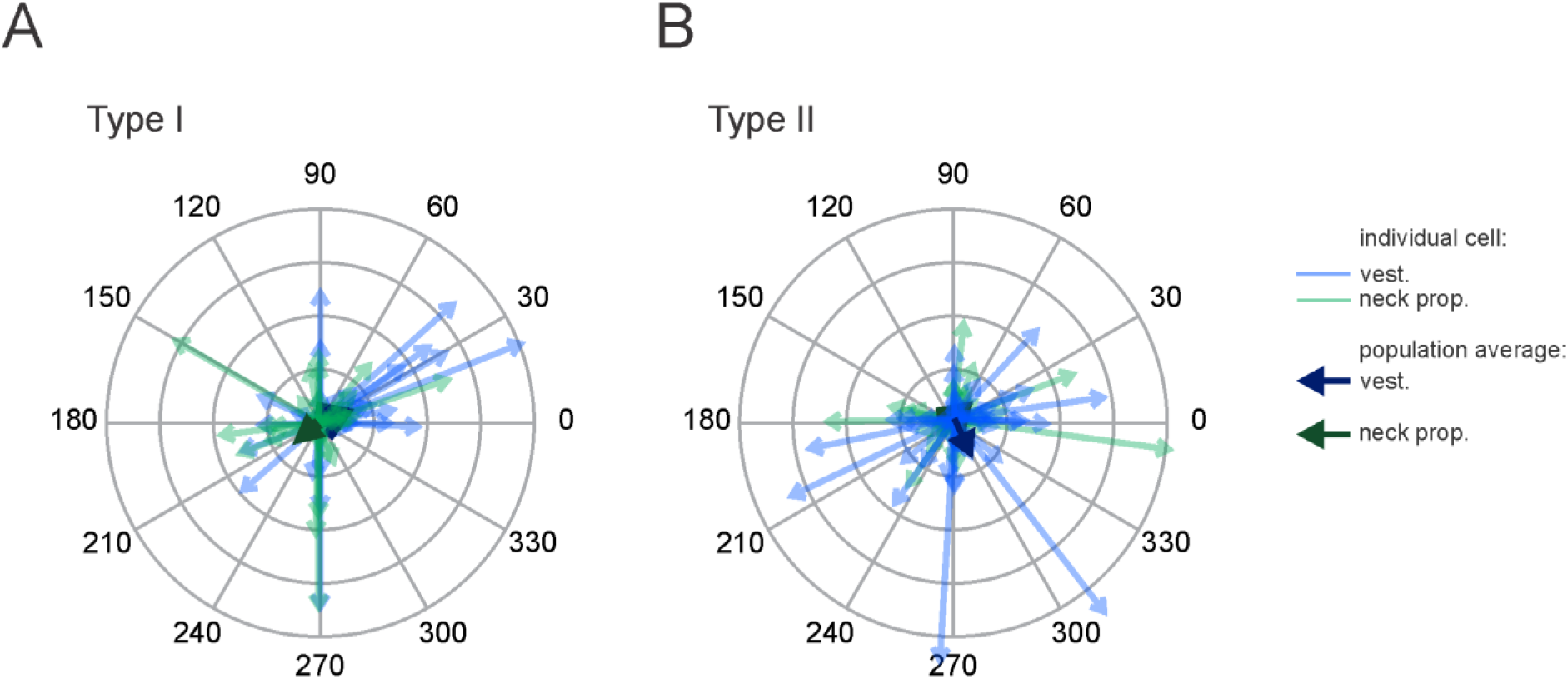
Purkinje cell responses to combined vestibular–neck proprioceptive stimulation in the non-preferred direction of vestibular stimulation. **(A, B)** Polar plots illustrating the vestibular (blue) and neck proprioceptive (green) neuronal response sensitivities of Type I **(A)** and Type II **(B)** Purkinje cells for non-preferred direction of vestibular stimulation and complementary direction proprioceptive stimulation (i.e., body-under-head motion). Superimposed blue and green arrows represent the mean population vectors, respectively.

**Figure 4 - figure supplement 1.**
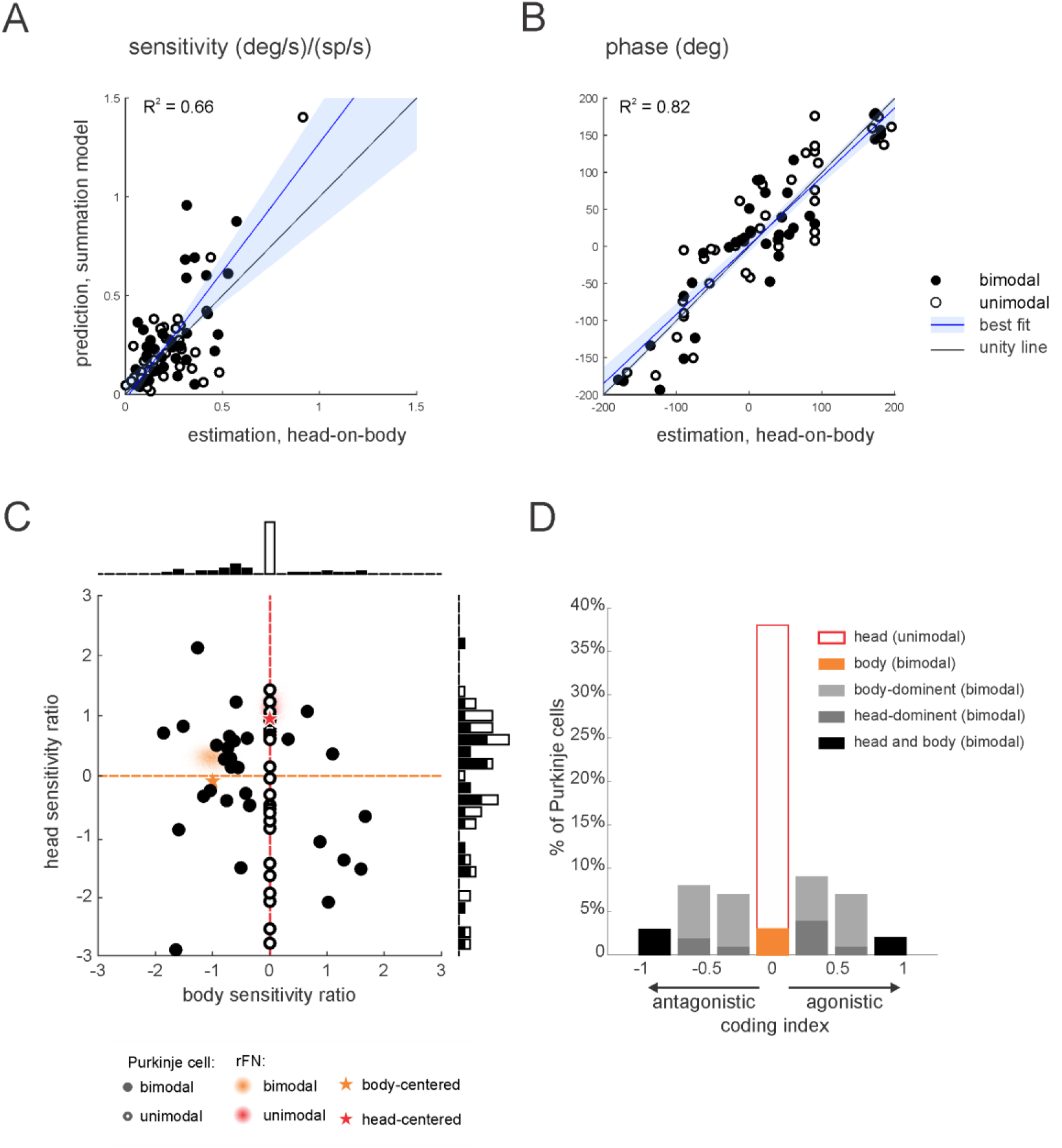
Purkinje cell simple spike responses to combined stimulation are well predicted by the linear summation of a given neuron’s responses to vestibular and proprioceptive stimulation when applied alone. **(A,B)** Comparison of estimated and predicted sensitivities (**A**) and phases (**B**) of Purkinje cell responses to head-on-body rotations in the non-preferred movement direction. The linear summation of a given neuron’s vestibular and neck proprioceptive sensitivities well predicted both measures in the combined condition. Blue lines and shading denote the mean ± 95% CI of linear fit. **(C)** Scatter plot of the relationship between the head sensitivity ratio (S_vest._+prop. / S_vest._) and body sensitivity ratio (S_prop._ / S_vest._). Histograms (top and right) illustrate the distributions of body and head sensitivity ratios, respectively. Orange versus red stars indicate ideal neurons encoding body versus head movement in space, respectively. For comparison, the red and orange shaded areas representing the distribution of values estimated for unimodal and bimodal rFN neurons (Brooks et al. 2009) are superimposed. **(D)** Distribution of coding indexes (see Methods). Positive and negative values correspond to agonistic and antagonistic responses to head vs. body encoding, respectively.

**Figure 6 - figure supplement 1.**
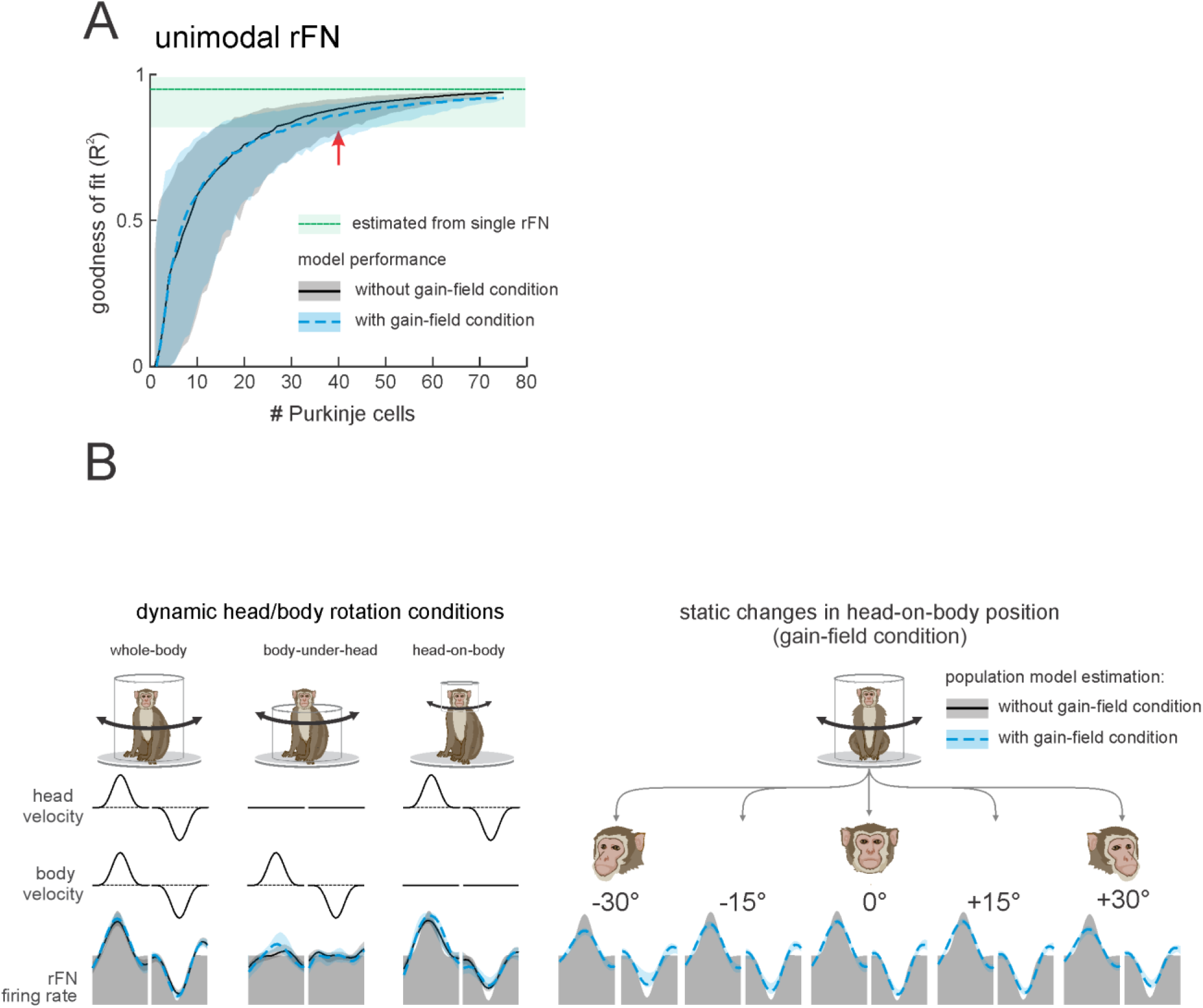
A simple linear population model of Purkinje cell integration can explain the responses of target unimodal neurons in deep cerebellar nuclei across all self-movement conditions. **(A)** Model performance as a function of the number of Purkinje cells. Black curves correspond to the population modeling that was performed on the simple spike firing rate of all 73 Purkinje cells recorded during three passive conditions (i.e., whole-body, body-under-head, and head-on-body movements. Blue curves correspond to the modeling of the simple spike firing rate of all 73 Purkinje cells during three passive conditions as well as the simulated responses of these cells in the static head-on-body position condition (**Fig. 5**). The variability estimated from population of rFN unimodal neurons previously described by Brook and Cullen (2009) is represented by the green shaded band. **(B)** Typical firing rate of unimodal neurons in the rostral fastigial nucleus (rFN) (grey shaded region) and the estimated firing rate based on a population of 50 Purkinje cells (mean ± SD) during three dynamic head/body rotation conditions (left) and the static changes in head-on-body position condition (right). We also performed population modeling on rFN type II cell and found similar results.

## Notes

### Competing Interest Statement

The authors have declared no competing interest.

